# Using a combination of CRISPR/Cas9, behavioural experiments and functional analysis to characterise taste receptors in honeybees

**DOI:** 10.1101/2022.03.17.484777

**Authors:** Laura Değirmenci, Fábio Luiz Rogé Ferreira, Adrian Vukosavljevic, Cornelia Heindl, Alexander Keller, Dietmar Geiger, Ricarda Scheiner

**Author notes:** L.D. and F.L.R.F. contributed equally to this work.

## Abstract

Honeybees (*Apis mellifera*) need their fine sense of taste to evaluate nectar and pollen sources. Gustatory receptors (Grs) translate taste signals into electrical responses. *In vivo* experiments primarily demonstrate collective responses of the whole Gr-set, but little is known about the individual impact of receptors. Here, we disentangle for the first time the contributions of three gustatory receptors (AmGr1-3) in sugar sensing of honeybees by combining CRISPR/Cas9 mediated genetic knock-out, electrophysiology and behaviour. AmGr1 responds to multiple sugars. Bees lacking this receptor have a reduced response to sucrose and glucose but not to fructose. AmGr2 acts as co-receptor of AmGr1 in a heterologous expression system, but honeybee knock-out mutants perform normally. Eliminating AmGr3 while preserving AmGr1 and AmGr2 abolishes the perception of fructose but not of sucrose. We thus dissociate the roles of AmGr1, AmGr2 and AmGr3 in honeybee taste perception.

## Introduction

Honeybees are important pollinators of crops and wild plants world-wide [1]. They depend on floral nectars as their main source of carbohydrates, which are converted into honey stores for provisioning the colony. It is therefore essential for honeybees to perceive and evaluate sweetness. Surprisingly, honeybees only possess a very reduced set of ten gustatory receptors (Grs) in their genome [2] [3] compared to the genome of other insects like the fruit fly (*Drosophila melanogaster*) with 69 genes or that of the mosquito (*Anopheles gambiae*) with up to 75 genes coding for gustatory receptors [2]. Despite this small set of Grs, honeybees can perceive many different sugars including sucrose, fructose, glucose, maltose, trehalose and melezitose. Most of these occur in nectar or honeydew in varying concentrations, while trehalose functions as main blood sugar in bees and other insects [4] [5] [6] [7] [8] [9] [10].

The sugar receptors of the honeybee are located on their antennal tips, their front tarsi and their mouthparts, but also internally in brain and gut [3] [11]. While AmGr3 is a specific fructose receptor [12] [13], AmGr1 has been shown to detect a broad spectrum of sugars including sucrose, glucose and trehalose in a heterologous expression system [14]. So far, AmGr2 has been assumed to act solely as a co-receptor of AmGr1, when studied in *Xenopus* oocytes [12] [13] [14]. Based on the taste range of honeybees and the structural similarity of many sugar molecules, AmGr1 and AmGr2 (as its co-receptor) should also respond to other sugars. However, many details of their function and potential interaction are still lacking. The new genome-editing technique of CRISPR/Cas9 knock-out, which has been employed successfully in numerous insects such as *Drosophila melanogaster* [15], *Aedes aeqypti* [16], *Bombyx mori* [17] and recently in the honeybee [13] together with a refined electrophysiological analysis of gustatory receptors heterologously expressed in *Xenopus* oocytes allow us to separate the individual roles of Grs and answer important open questions regarding their impact on taste perception in this model insect.

## Results

### AmGr1 represents an essential component for the sucrose and glucose taste *in vivo* and the heterologous expression system of *Xenopus* oocytes

Our TEVC measurements confirm AmGr1 as a receptor for sucrose and glucose (as shown previously in [14]), that does not detect fructose, when transiently expressed in *Xenopus* oocytes (Fig. 1 A1). Oocytes co-expressing all three sugar receptors (AmGr1-3, representing a wildtype-like set of sugar receptors) elicited sustained inward currents of several nano amps when flushed with sucrose and fructose, whereas currents elicited during glucose application displayed a differed trace shape (Fig. 1 A2). Although glucose was present for 60 s, the glucose-induced currents appeared only transiently for about 20 s (see also below).

**Figure 1:**
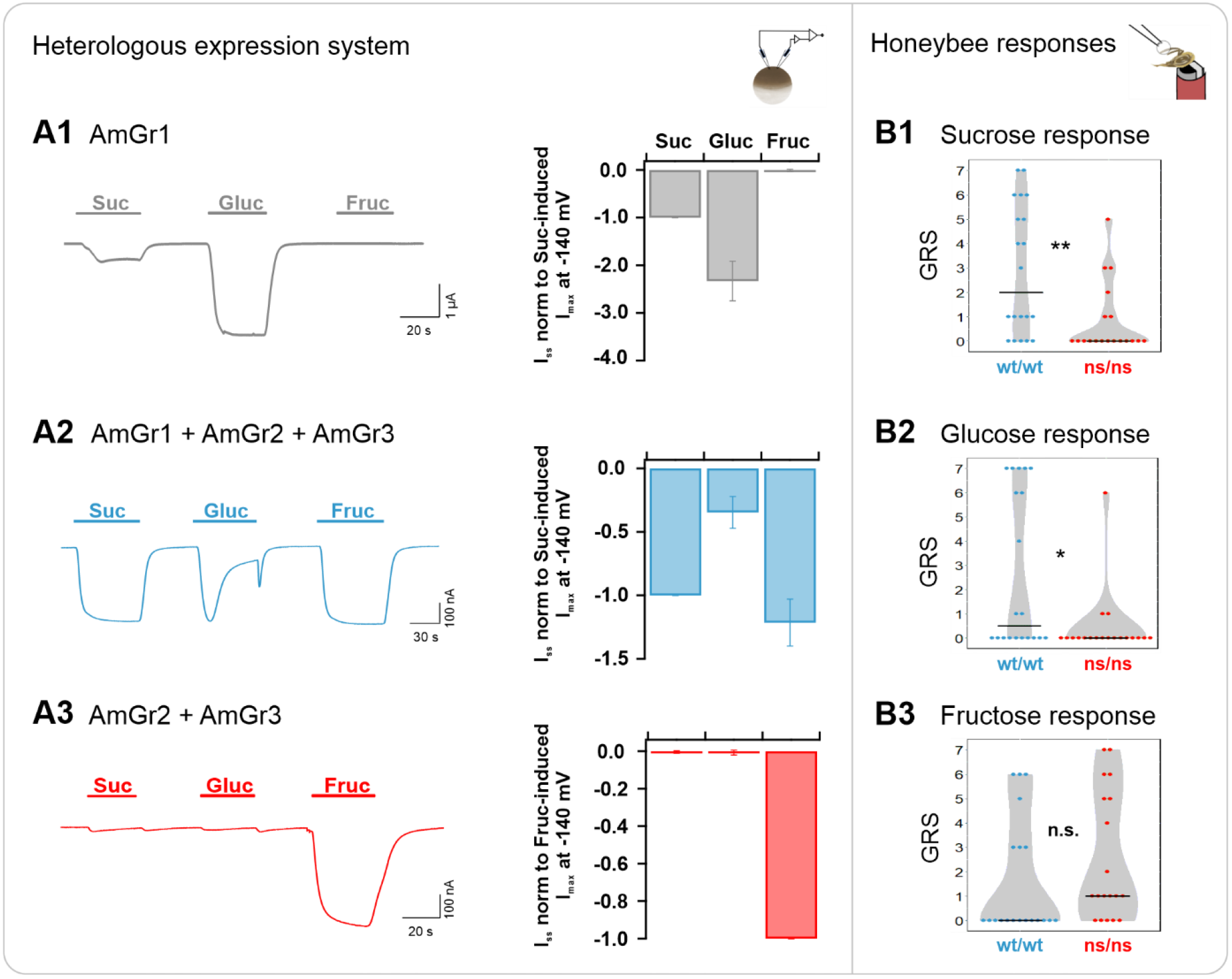
Functional analysis of *A. mellifera* gustatory receptor AmGr1 using a matched heterologous expression system and *in vivo* comparative approach. **A1-3: two-electrode voltage clamp measurements:** Current traces were recorded at a holding potential of −80 mV in response to perfusion with sucrose (Suc), glucose (Gluc) and fructose (Fruc) in standard solution (left panel). Sugar-induced steady-state currents (*I_SS_)* were recorded at a membrane potential of −140 mV (middle panel). **(A1)** Representative whole oocyte current trace of AmGr1-expressing oocyte (left panel); Quantification of sugar-induced *I_SS_.* Currents were normalized to sucrose-evoked *I_SS_* at −140 mV (mean of *n* = 16 oocytes ± SD; middle panel). **(A2)** Inward whole oocyte currents from AmGr1/AmGr2/AmGr3-expressing oocyte (wild-type mimicry; left panel); Quantification of sugar-induced *I_SS_* that were normalized to sucrose-evoked *I_SS_* at −140 mV (mean of *n* = 10 oocytes ± SD, middle panel). **(A3)** Whole oocyte currents from AmGr2/AmGr3 co-expressing oocyte (left panel); Quantification of sugar-induced *I_SS_.* Currents are normalized to the fructose-evoked *I_SS_* at −140 mV (mean of *n* = 9 oocytes ± SD; middle panel). **B1-3: behavioural evaluation through proboscis extension response (PER, *in vivo*).** Wild-type (wt/wt, N=20) and AmGr1 mutant bees (ns/ns; N=19) were presented a series of sugar concentrations (16%, 20%, 25%, 32%, 40%, 50% and 60%) of all three sugars sucrose **(B1)**, glucose **(B2)** and fructose **(B3)**. The sum of the responses (PERs) towards the concentrations of one of the sugars was recorded as a sugar-specific GRS (gustatory response score). AmGr1 mutants were less responsive to sucrose when compared with wild-type bees and had significantly lower GRS (B1; Mann-Whitney-U, ns/ns vs. wt/wt, p=0.0032, **). Glucose responsiveness in AmGr1 mutants was significantly lower than that in wild-types (B2; Mann-Whitney-U, ns/ns vs. wt/wt, p=0.0125, *). Both groups did not differ in fructose GRS (B3; Mann-Whitney-U, ns/ns vs. wt/wt, p=0.0779, n.s.).

To verify the impact of AmGr1 on overall sugar response, co-expression of only AmGr2 and AmGr3 in oocytes was tested, simulating a honeybee mutant lacking functional AmGr1 receptor. Under this scenario, only fructose-induced macroscopic currents could be observed (Fig. 1 A3), indicating that AmGr2 does not appear to act as a co-receptor of the fructose-specific receptor AmGr3 [12] [13] in *Xenopus* oocytes.

Testing sugar responsiveness at the behavioural level using the established proboscis extension response assay revealed that AmGr1 honeybee mutants were significantly less responsive to sucrose than wild type bees when the sugar was applied to their antennae (Fig. 1 B1). The mutants also displayed a reduced responsiveness to glucose (Fig. 1 B2) compared to wildtype bees. However, the fructose responses of AmGr1 mutants did not differ from those of wild type bees (Fig. 1 B3).

### AmGr2 is a co-receptor for sucrose and glucose perception

Although expression of AmGr2 in oocytes could be confirmed using a N-terminal fused YFP as a genetically encoded reporter protein (Fig. S2 A), AmGr2-expressing oocytes did not reveal any macroscopic sugar-induced currents in TEVC experiments (Fig. 2 A1). However, reproducible microscopic inward current deflections in response to sugar application could be recorded – albeit very low – reaching current amplitudes as few as tens of nano amps overall (Fig. 2 A1, inset). Intriguingly, sustained inward currents could only be generated by application of sucrose but not with any of the other sugars. Analogous to the glucose-induced current responses in the wildtype simulation in oocytes (Fig. 2 A2), AmGr2-expressing oocytes revealed transient inward currents during glucose application followed by a rapid remission to the baseline. Its peak amplitude reached the current values in the same range as recorded for sucrose. A similar transient current deflection was observed upon washout with a reference solution. Perfusion with fructose evoked a comparable current response pattern to AmGr3 expressing oocytes either alone or in combination with AmGr1 and/or AmGr2. Furthermore, bimolecular fluorescence complementation (BiFC) experiments confirmed physical interaction of AmGr2 subunits, indicating the assembly of homomeric AmGr2 receptors of low electric activity (Fig. S2 B). Taken together, the sugar-induced inward currents, along with the physical interaction between AmGr2 subunits proven with BiFC, suggest that AmGr2 is able to assemble to a functional homomeric channel that builds up an ion pore, thus being able to perform ligand-gated channel activity with low conductance in oocytes by itself.

**Figure 2:**
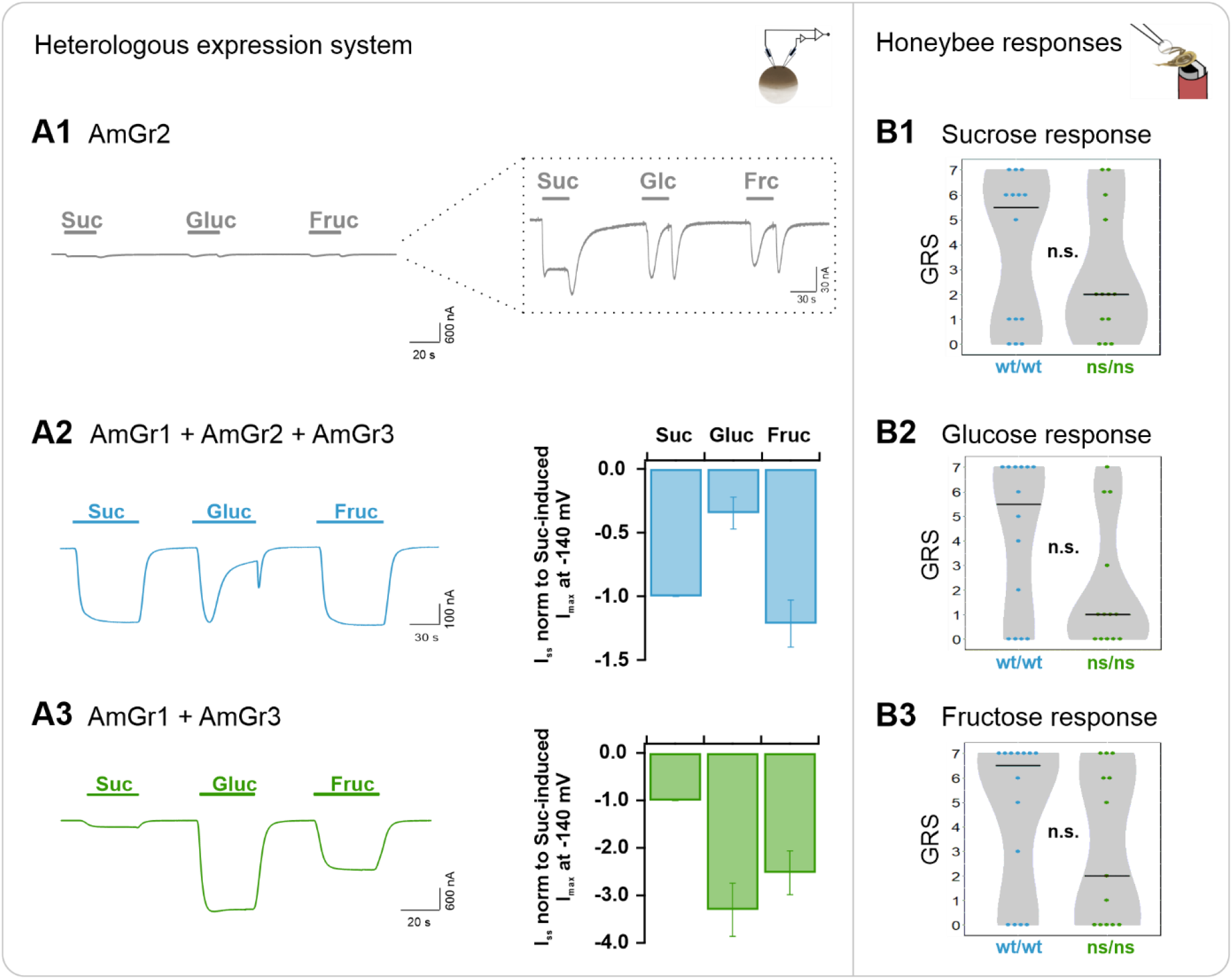
Functional analysis of *A. mellifera* gustatory receptor AmGr2 using a matched heterologous expression system and *in vivo* comparative approach. **A1-3: two-electrode voltage clamp measurements:** Current traces were recorded at a holding potential of −80 mV in response to perfusion with sucrose (Suc), glucose (Gluc) and fructose (Fruc) in standard solution (left panel). Sugar-induced *I_SS_* were recorded at a membrane potential of −140 mV and normalized to the currents in either sucrose or fructose solution (middle panel). **(A1)** Representative whole oocyte current trace of AmGr2-expressing oocyte. Inset: Magnification of the current trace reveals microscopic sustained or transient inward currents upon sugar application. **(A2)** Inward whole oocyte currents from AmGr1/AmGr2/AmGr3-expressing oocyte (wild type mimicry, left panel); Quantification of sugar-induced *I_SS_* that were normalized to sucrose-evoked *I_SS_* at −140 mV (mean of *n* = 10 oocytes ± SD; middle panel). **(A3)** Whole oocyte currents from AmGr1/AmGr3 co-expressing oocyte (left panel); Currents are normalized to the fructose-evoked *I_SS_* at −140 mV (mean of *n* = 13 oocytes ± SD; middle panel). **B1-3: behavioural evaluation through proboscis extension response (PER, *in vivo*).** Wild-type (wt/wt, N=14) and AmGr2 mutant bees (ns/ns; N=13) were presented a series of sugar concentrations (16%, 20%, 25%, 32%, 40%, 50% and 60%) of all three sugars sucrose **(B1)**, glucose **(B2)** and fructose **(B3)**. The sum of the responses (PERs) towards the concentrations of one of the sugars was recorded as a sugar-specific GRS (gustatory response score) of each respective bee. AmGr2 mutants (ns/ns) did not show any significant differences in their responsiveness towards all three sugars when compared to wild-type (wt/wt) bees, neither to sucrose (B1; Mann-Whitney-U, ns/ns vs. wt/wt, p=0.5351, n.s.), to glucose (B2; Mann-Whitney-U, ns/ns vs. wt/wt, p=0.0909, n.s.) or to fructose (B3; Mann-Whitney-U, ns/ns vs. wt/wt, p=0.2536, n.s.).

Sucrose and glucose stimulation of oocytes co-expressing solely AmGr1 and AmGr3, lacking the expression of AmGr2, led to a very similar current response pattern as that seen for the sole expression of AmGr1, with additional fructose-induced currents upon fructose application (Fig. 2 A3). Honeybee AmGr2-mutants did not differ from wild-type bees in their responses to sucrose, glucose or fructose when tested in the behavioural assay (Fig. 2 B1, B2 and B3). These findings indicate that AmGr2 on its own does not provide sufficient ion channel performance on a comparable scale such as sugar receptors AmGr1 nor AmGr3 but can unfold its function in honeybee sugar sensation acting as a co-receptor for AmGr1.

### AmGr3 is a specific fructose receptor

The sole expression of AmGr3 in oocytes only led to fructose-induced current responses (Fig. 3 A1). Neither sucrose nor glucose acted as ligands of AmGr3 in our experiments, supporting earlier findings [12] [13]. Oocytes co-expressing AmGr1 and AmGr2, simulating the honeybee AmGr3 knock-out mutant, did not elicit any fructose-induced currents in the cells (Fig. 3 A3). Honeybee AmGr3 homozygous mutants displayed a significantly reduced response to fructose compared to wild-type bees, what was found out and published previously [13]. This difference was not observed when tested with sucrose in the PER paradigm. These findings show that AmGr3 is unequivocally a fructose-specific receptor in the honeybee demonstrated by heterologous expression system and behaviour *in vivo*.

**Figure 3:**
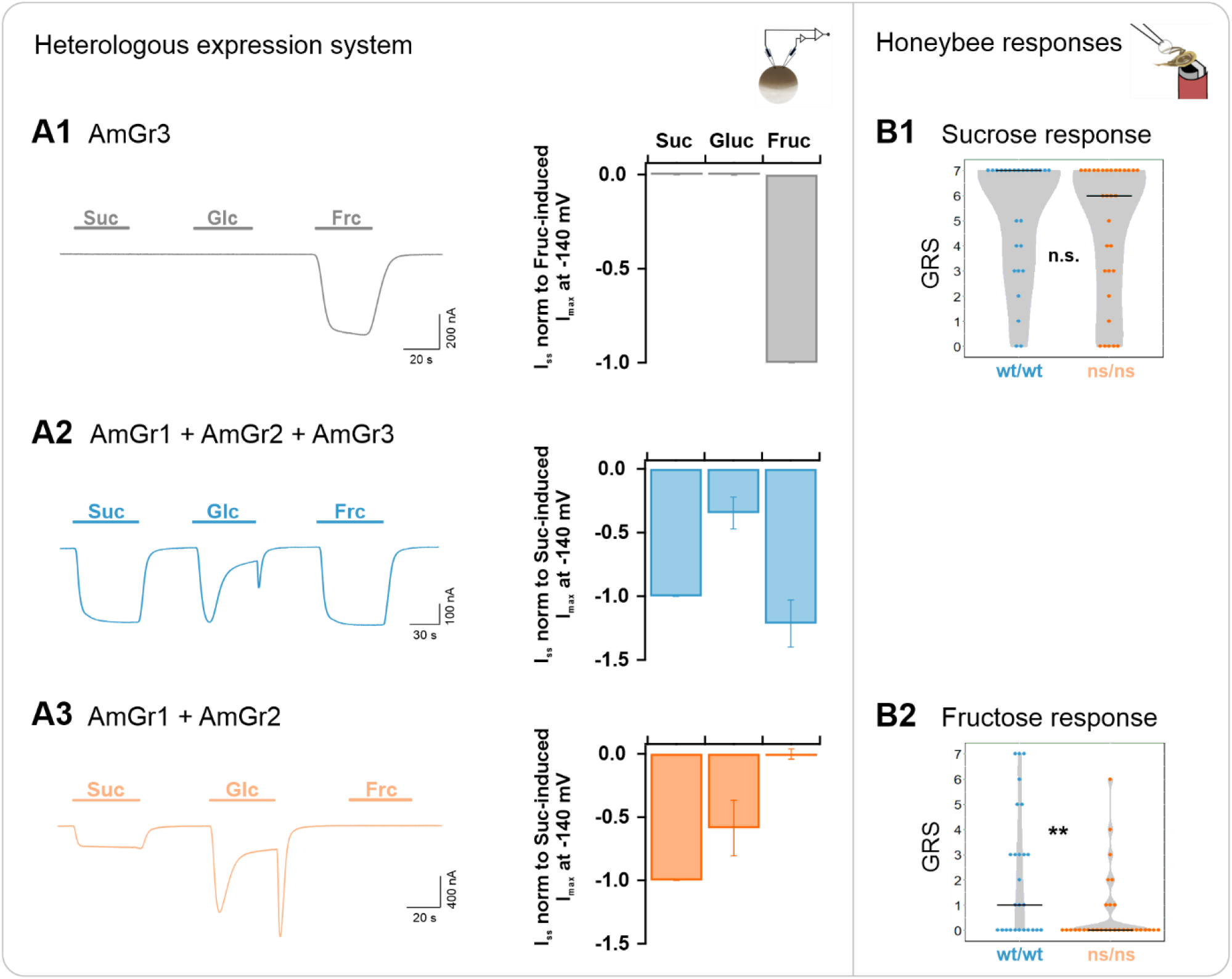
Functional analysis of *A. mellifera* gustatory receptor AmGr3 using a matched heterologous expression system and *in vivo* comparative approach. **A1-3: two-electrode voltage clamp measurements:** Current traces were recorded at a holding potential of −80 mV in response to perfusion with sucrose (Suc), glucose (Gluc) and fructose (Fruc) in standard solution (left panels). Sugar-induced *I_SS_* were recorded at a membrane potential of −140 mV (middle panels). **(A1)** Representative whole oocyte current trace of AmGr3-expressing oocyte (left panel); Currents are normalized to the fructose-evoked *I_SS_* at −140 mV (mean of *n* = 8 oocytes ± SD; middle panel). **(A2)** Inward whole oocyte currents from AmGr1/AmGr2/AmGr3-expressing oocyte (wild-type mimicry, left panel); currents were normalized to sucrose-induced *I_SS_* at −140 mV (mean of *n* = 10 oocytes ± SD; middle panel). **(A3)** Whole oocyte currents from AmGr1/AmGr2 co-expressing oocyte (, left panel); currents were normalized to sucrose-induced *I_SS_* at −140 mV (mean of *n* = 13 oocytes ± SD; middle panel). **B1-2 (as published previously in Degirmenci et al. 2020): behavioural evaluation through proboscis extension response (PER, *in vivo*).** Wild-type (wt/wt, N=26) and AmGr3 mutant bees (ns/ns; N=31) were presented a series of sugar concentrations (16%, 20%, 25%, 32%, 40%, 50% and 60%) of the sugars sucrose **(B1)** and fructose **(B2)**. The sum of the responses (PERs) towards one of the sugars was recorded as a sugar-specific GRS (gustatory response score). AmGr3 mutants did not show any difference in GRS when compared with wild-type bees (B1; Mann-Whitney-U, ns/ns vs. wt/wt, p=0.4279, n.s.). Fructose GRS of AmGr3 mutants were significantly lower than those of wild-types (B2; Mann-Whitney-U, ns/ns vs. wt/wt, p=0.0062, **).

### Modulation of sugar-induced signals by receptor co-expression

*Xenopus* oocytes expressing all three receptors showed robust current deflections in TEVC when exposed to sucrose, glucose or fructose. Interestingly, glucose-induced inward currents were of transient nature only, showing a decay over the course of application (Fig. 1 A2, Fig. 2 A2 or Fig. 3 A2). Following the decay, glucose-induced steady-state currents reached similar levels of maltose, trehalose and melezitose (Fig. S1 F, bar diagram). This behaviour is also apparent when AmGr3 is absent (Fig. 3 A3. Fig. S1 C). However, when AmGr1 was expressed alone, the glucose current trace was stronger and did not decay over time (Fig. 1 A1). In agreement with Jung et al. [14], AmGr2 itself did not show macroscopic sugar-induced currents (Fig. 2 A1) which is well in line with honeybee AmGr2 mutants that did not show significant differences in responses to sucrose, glucose or fructose compared to controls (Fig. 2 B1, B2 and B3). Moreover, our BiFC experiments indicate a direct physical interaction between AmGr1 and AmGr2, strongly suggesting that heteromerization occurs (Fig. S2 B). Thus, AmGr2 seems to act exclusively as co-receptor for the sucrose signal of AmGr1. AmGr2 additionally modulates the strength [14] and the time-dependent characteristics of AmGr1-derived glucose signals.

For fructose induced currents – caused by AmGr3-expression as described above – no obvious modulations could be found when it was co-expressed with the other receptors (Fig. 1 A3 or Fig. 2 A3). Mutant bees lacking AmGr2 (but expressing AmGr3 and AmGr1) as well as AmGr1 homozygous mutants (with AmGr3 and AmGr2 being present) did not show any significant difference in their fructose response compared to wildtype bees (Fig. 1 B3 and Fig. 2 B3). Thus, among the tested gustatory receptors, AmGr3 is irreplaceable as the specific fructose receptor [13]. Moreover, its electrophysiological properties are not modulated by any of the other sugar receptors.

## Discussion

Taste plays a critical role for honeybees when evaluating and choosing profitable food sources in terms of concentration and type of sugar [28]. Bees rely on pollen as a source of protein and nectar or honeydew as their main sources of carbohydrates. It has been shown that honeybees perceive e.g. the sugars sucrose, glucose, fructose, maltose and melezitose, which are the main constituents found in their food [5] [6] [28]. It is therefore astonishing that honeybees only have a reduced set of gustatory receptors and, among them, only three are sugar receptors (AmGr1-3) [2] [3]. Bees show a broad spectrum of food-related and taste-related behaviours, like division of labour [29] [25] or the ability to memorize sugars and associate odours with them [30]. Differences in sugar receptor expression were discovered as a possible regulatory mechanism of division of labour [3] [31].

To elucidate the complexity of sugar perception in honeybees, we carried out a combined study on individual sugar receptors using the oocyte heterologous expression system and homozygous receptor mutants produced with CIRPSR/Cas9 *in vivo*. While our functional analysis illustrates the initiating step of taste perception on a molecular level (sugar-induced receptor activation), our *in vivo* analysis focusses on the behavioural outcome (PER index) as consequence. In agreement with Jung et al. [14], the gustatory receptor AmGr1 used in our experiments responded to sucrose and glucose (as well as to maltose, trehalose and melezitose; data shown in suppl. Fig. S1). At the behavioural level, this receptor is involved in the evaluation of sucrose, glucose and maltose (data shown in suppl. Fig. S2), because bees lacking AmGr1 were less responsive compared to controls. The promiscuous ligand specificity of AmGr1 suggests that the taste perception system of honeybees counterbalances a limited set of Grs. Interestingly, all sugars stimulating AmGr1 comprise at least one accessible D-glucose unit and most of these sugars play an important role in the bee's ecological context, i.e. in their food sources (sucrose, fructose, glucose, maltose and melezitose) or their blood sugar (trehalose). In contrast, we could not detect any responses to the less relevant sugars such as arabinose, mannose or galactose and none of them contains a glucose unit. Additionally, no signal was generated by raffinose application. We therefore assume that the glucose unit in raffinose is embedded and difficult to access. The intramolecular arrangement of the glucose units seems to play a critical role in the sugar recognition by AmGr1. Future experiments combining structure-related functional analysis with glucose analogues might provide new insights regarding its sugar stereospecificity.

The gustatory receptor AmGr2 acts as a co-receptor modulating the specificity of AmGr1 but not of AmGr3. AmGr2 co-expression tunes the broad sugar receptor (AmGr1) into a sucrose receptor, preserving sucrose recognition but drastically affecting glucose signals (currents displayed a transient response only). This inhibition-like property of AmGr2 could be related to a desensitization effect, partly influenced by substrate concentration. A similar desensitization effect has been described for the human epithelial sodium channel ENaC, which is thought to interact with environmental cues, such as ionic or acidic concentrations [32] [33] and which undergoes self-inhibition, modulated by extracellular sodium concentration [34]. For instance, cryo-EM 3D structures of conformations in high pH resting and low pH desensitized conformations of the splice variant ASIC1a from Acid-Sensing Ion Channels (ASICs, as proton-gated members of the ENaC superfamily) show ion permeation and proton-dependent gating via a peptide loop at the N-terminus [35]. Even though these channels are structurally unrelated to insect gustatory receptors and information about structural domains within Grs throughout the *Insecta* taxon are scarce, these examples display how molecular gating mechanisms can be relevant in receptor modulation. Assembly of AmGr1 and the co-receptor AmGr2 into heterotetramers seem to adopt an altered, yet fine molecular gating mechanism that restricts the ion passage when interacting with ligands other than sucrose. These desensitization-like kinetics could be interpreted as a hint towards substrate-induced inhibition during the open-state channel gating. Therefore, a broad spectrum of sugar taste in honeybees does not necessarily have to be constant but can rather be fine-tuned to a small set of sugars. Thus, heteromerization between AmGr1 and AmGr2 broadens their functional diversity [36] [37]. Detailed structural information and experiments with site-directed mutagenesis of putative motifs are indispensable to provide insights into the mechanistic contribution of motifs of individual subunits, which might deliver a basis for understanding Gr modulation and their possible role in desensitization.

AmGr2 knock-out mutants did not show any significant changes in behaviour in response to the sugars sucrose, glucose and fructose. Though the PER paradigm works very well to fully elucidate receptor activity [13], it might be rather unspecific for the co-receptor AmGr2. For instance, for the fruit fly (which inheres 68 gustatory receptors [2]), it was shown that the neurons are widely distributed throughout the main gustatory organs, each expressing distinctive types and sets of Grs. These heterogeneous Gr populations can be found in the same organ or sensilla [38], they might respond to different sugars [39] and even interact with each other [40]. Assuming a comparable complexity in the honeybee as well as further processing in the brain, the actual influence of the co-receptor AmGr2 in the behavioural outcome might be blurred. This could also be the reason for our finding that an absolute loss of response could not be observed in all mutants [36]. Fixation could also influence natural behaviour [41]. Earlier experiments showed that freely moving bees or bees in cages prefer sucrose over all other sugars [4], [42] comparable to fixed bees in more recent PER experiments [43]. In all TEVC experiments, however, the sucrose signals measured were weaker or similar to those of glucose. The yet uncovered co-receptor function of AmGr2 or differences in general receptor expression might be factors modulating the receptor signal and the actual behaviour, but these points need further investigation.

Our results clearly show that AmGr3 is a specific fructose receptor. Inward currents induced by this sugar can no longer be measured when AmGr3 is missing in the cell system and the homozygous honeybee mutants are significantly less responsive to fructose. The fructose signal does not seem to be influenced by any other receptor (such as AmGr1 by AmGr2). The homolog of AmGr3 in *Drosophila* appears to be a sensor for saturation of these insects, displayed by the fructose level, which seems to react to food intake regardless of actual blood sugar [44] [45]. The specificity of AmGr3 might indicate a comparable function in honeybee saturation with sugars.

Overall, our matched *in vivo* and functional analysis with homozygous mutants as well as the biophysical analysis in *Xenopus* oocytes was successful in providing a link between the electrophysiological response and the behavioural response to sugars in honeybees. Thus, the repertoire of sugar taste in honeybees could be mimicked in *Xenopus* oocytes and used to further uncover open questions regarding sugar perception in honeybees. Since individual homozygous mutants survived to adulthood, we demonstrated that all sugar receptors (AmGr1-3) are not essential for larval development in honeybees. Homozygous mutants generated with CRISR/Cas9 can complement studies in live animals, assuming that the examined receptors have direct ligands and are linked to relevant behaviours such as was shown for the main sugar receptor AmGr1 and the fructose receptor AmGr3. For the AmGr2 co-receptor we show in the cell system that it disrupts sustained glucose signals of AmGr1 compared to sucrose signals. Similar to how complex colour vision can be achieved by just three photoreceptors (trichromatic vision) [46], our investigations show that the broad sugar taste in honeybees is regulated in a complex way and can be covered by only three sugar receptors.

## Material and Methods

### *Xenopus* oocytes preparation

TEVC experiments were performed in oocytes from mature female *Xenopus laevis* (African clawed) frogs. They were healthy, non-immunized and have not been involved in any previous procedures. The Julius-von-Sachs Institute of Wuerzburg University holds the permission to keep these animals and is registered at the government of Lower Franconia, ref. No. 55.2-2532-2-1035. The conditions for keeping the frogs were at 20 °C temperature, with a 12/12 h day/night cycle, in dark grey 96 l tanks (5 frogs/tank) covered with a see-through aluminium grid. Feeding with floating trout food (Fisch-FitMast 45/7 2 mm, Interquell GmbH, Wehringen, Germany) took place in a fixed bi-weekly schedule. As hiding spots for the frogs, 30 cm long PVC pipes with a diameter of around 10 cm were placed in the tanks. Aquarium pumps ensured a continuous water circulation and filtration. Oocytes were obtained through surgical removal of ovary lobes (partial ovariectomy). The female *X. laevis* frogs were anesthetized by immersion in water containing 0.1 % 3-aminobenzoic acid ethyl ester ca. 0.5 h prior to the operation. Subsequently, oocytes were partially freed from the ovary membrane and treated with collagenase I in Ca^2+^-free ND96 buffer (10 mM HEPES pH 7.4, 96 mM NaCl, 2 mM KCl, 1 mM MgCl_2_) for 1-1.5 h. Afterwards, oocytes were washed with Ca^2+^-free ND96 buffer and kept at 16°C in ND96 solution (10 mM HEPES pH 7.4, 96 mM NaCl, 2 mM KCl, 1 mM MgCl_2_, 1 mM CaCl_2_) containing 50 mg/l gentamycin. For electrophysiological experiments, stage V or VI oocytes were injected with cRNA and incubated for 2-5 days before TEVC measurements were performed. The final cRNA concentration used for injection was adjusted to fixed amounts of 25 ng AmGr1, 50 ng AmGr2 and 50 ng AmGr3, independent of sole expression or co-expression combinations.

### RNA extraction and cDNA synthesis

To extract RNA, frozen honeybee tissue (antennae, mouthparts and tarsi) was broken up in 750 μl (TriFast USA) in a 2 ml Eppendorf (Hamburg, Germany) tube using the TissueLyzer (QIAGEN, Venlo, Netherlands) with stainless steel beads (5 mm). The sample was allowed to incubate for 5 min before 200 μl chloroform was added. Then it was mixed and centrifuged. The resulting aqueous phase was transferred to a Perfect Bind RNA Colum of the Total RNA Kit (peqGOLD, VWR, Radnor, USA), following the kit’s protocol for purification. Precipitation of RNA was performed with 3 M sodium acetate, washed with ethanol, dried and adjusted to the required concentration when resolved. The AccuScript Hi-Fi Kit and Oligo(dT) primers (18mers) were used for cDNA synthesis following the manufacturer’s instructions and using concentrations required (Agilent Technologies, Santa Clara, USA). An RNA digestion step was added using the enzyme RNAse H (NEB, Ipswich, USA) according to the protocol. Following the instructions, large scale Phusion PCRs were performed (see suppl. Tab. S1).

PCR products were run on a 1 % (w/v) agarose gel and the appropriate bands were cut out and purified according to the Wizard SV Gel and PCR Clean-Up System (Promega, Fitchburg, USA). A 20 min A-tailing step (72 °C, 0.2 mM dATP) was performed with taq polymerase (NEB, Ipswich, USA).

### Cloning and cRNA synthesis

The prepared PCR fragments were inserted into the pGEM-T vector via T/A cloning following the manufacturer’s recommendations (Promega, Fitchburg, USA). Competent *Escherichia coli* were transfected with the ligation mix and plated on agar plates as described previously [13]. *E. coli* could grow over night at 37 °C, clones were picked with the blue-white selection and transferred to an overnight culture (as described in detail in [13]). After pelleting, the isolation was performed according to the Plasmid Miniprep Kit I (peqGOLD, VWR, Radnor, USA) and verified through sequencing.

The complementary DNA (cDNA) of each receptor gene (AmGr1-AmGr3) was sub-cloned into pNBIu (based on pGEM vectors) oocyte expression, YFP-fusion and BiFC (Bimolecular Fluorescence Complementation) vectors using an advanced uracil-excision-based cloning technique in combination with a PfuX7 polymerase, as described by Nour-Eldin et al. and Nørholm et al. [18] [19]. Both the YFP and the complementary halves of YFP (for BiFC experiments see supplement) were cloned upstream of the respective AmGr-receptor cDNA as illustrated in suppl. Fig. S2. The construct inserts were verified by sequencing. Complementary RNA (cRNA) was prepared with the AmpliCap-Max T7 High Yield Message Maker Kit (Cellscript, Madison, WI, USA) according to the manufacturer’s specifications. For TEVC and BiFC experiments, oocytes were injected with 0.5 ng cRNA of each (co-expressed) construct.

### Oocyte recordings

To characterize the honeybee gustatory receptors (AmGr1-3), they were expressed either separately or co-expressed in different combinations in *Xenopus* oocytes and functional analysis was performed using the two-electrode voltage-clamp (TEVC) technique.

For sugar specificity measurements, reference solution was exchanged by either 160 mM of sucrose, glucose, fructose, maltose, arabinose, mannose, galactose, raffinose, trehalose or melezitose (ensuring a constant osmolarity in the range of 240 mOsmol/L).

TEVC recordings and data analysis: steady-state current (*I_SS_*) recordings of AmGr-expressing oocytes were established by performing a standard voltage protocol: an initial holding potential (V_H_) of 0 mV was followed by subsequent 200 ms single voltage pulses applied in the range of +20 mV to −140 mV in 20 mV decrements. The sugar-induced currents were derived by subtracting the currents in the absence of sugar from the currents in the presence of sugar. For each oocyte, the sugar-induced steady-state currents (*I_SS_*) were normalized either to the maximal absolute sucrose- or fructose-induced currents at −140 mV (depending on the respective AmGr-ensemble). Single-pulse measurements were performed under a constant −80 mV holding potential.

#### Recording solution

TEVC measurements with oocytes were performed using Tris/MES-based buffers. The standard solution contained 30 mM NaCl, 10 mM Tris/Mes (pH 7.5), 1 mM CaCl_2_, 1 mM MgCl_2_, 2 mM KCl and either 160 mM D-sorbitol (reference solution) or 160 mM of the tested sugars.

### Bimolecular Fluorescence Complementation (BIFC) assay

For documentation of YFP- or BIFC-derived fluorescence oocytes were excited with an argon laser line of 514 nm and YFP fluorescence emission was monitored between 500 and 580 nm. Pictures were taken with a confocal laser scanning microscope (Leica TCS SP5; Leica Microsystems CMS GmbH) equipped with a Leica HCX IRAPO L25×/0.95W objective. Images show a quarter of an oocyte.

### Preparation of sgRNA

Target-sites for the sgRNAs (single guide RNAs) were found in the first exons of the ORFs of the respective genes (AmGr1-3, see suppl. Tab. S2). Target specific sequences were found via benchling (https://benchling.com, San Francisco, USA) following the criteria mentioned elsewhere [13]. For each target an overlapping phusion PCR (NEB, Ipswich, USA) was performed with a forward primer including the T7 promoter and the certain crRNA sequence (specific for each sgRNA, see suppl. Tab. S2) and a stable revers primer (tracrRNA sequence, [13]). PCR products were purified, checked and used as template for the sgRNA synthesis. Purification of sgRNA was performed with the MEGAclear Transcription Clean-Up Kit (Invitrogen, Carlsbad, USA) after the DNAse digestion. As described, the quantified and pre-tested sgRNAs were aliquoted and frozen in their favorable concentration (best hatching rates and mutation rates were determined in pre-tests, data is not shown). During the experiment, a fresh aliquot of sgRNA and Cas9 enzyme (Cas9 Nuclease, S. pyogenes, 20 μM; NEB, Ipswich, USA) was used for each day and stored on ice (find respective concentrations in suppl. Tab. S2).

### Honeybee egg harvest

Beehives had related and naturally inseminated queens of *Apis mellifera carnica*, which were maintained outdoors at Würzburg University and fed with ApiInvert/ApiFonda (Südzucker, Mannheim, Germany) if necessary. Queens were caged in the JENTER systems (Karl Jenter GmbH, Germany) for three days. They were forced to lay their eggs through a comb-like cell grid and onto removable JENTER plug-in cells. Overnight eggs were discarded. We performed two replicate experiments per gene. For each, the time-monitored eggs were microinjected 0–1.5 hours after deposition (with either sgRNA for AmG1, AmGr2 or AmGr3 and water controls), so that the mutational event occurred in the single-cellular state, leading to fully mutated embryos without mosaic patterns. Roth et al. recently showed that frequently both alleles were mutated (homozygous mutants) and that the entire bee was affected (absence of mosaicism) [20]. Mutations were controlled via next generation sequencing (NGS, after preselection via fluorescence length analysis, FLA). Additionally, Yu and Omholt demonstrated in 1999 that division of the honeybee zygote is completed after 120±6.9 minutes, confirming our choice of procedure [21].

### Microinjection of eggs and artificial rearing of honeybees

Following the protocol of [20], eggs were processed and injected in a 35°C climate chamber, using the same set-up, procedure and material described in our prior work [13] [22]. Each egg was injected with 400 pl volume (water or sgRNA/Cas9 as described above, also see suppl. Tab. S2). Eggs were maintained in a sulfuric atmosphere until a few hours before hatching (1 ml of 16 % sulfuric acid per liter of volume, separated from the rings by a grid) and monitored. Hatched larvae were removed carefully with a modified Chinese grafting tool and transferred to prepared Nicot-wells (NICOTPLAST, Maisod, France) containing larval food. Larval rearing was carried out as described earlier [13] based on the protocol of [22]. One wing was removed from the freshly hatched and fully dried bees. Individuals were marked with coloured number plates (Opalith queen-marking plates) using super glue (UHU GmbH & Co. KG, Germany). All marked bees of one replicate (treated with sgRNA/Cas9 and the water controls) were placed into a cage with pollen and sugar water and maintained in an incubator at 35°C.

### Testing responsiveness to sugars

One-week-old bees were tested for their proboscis extension response (PER) to increasing concentrations of sucrose, glucose and fructose (in two replicates per gene) according to [13]. Prior to the test, each bee was immobilized on ice, carefully mounted in brass tubes and fixed with adhesive tape [23]. Subsequently, both antennae were stimulated with a droplet of a certain sugar water concentration. All three sugars (two for AmGr3), alternatingly starting with sucrose, fructose or glucose, were tested. After the water pre-test, an increasing concentration series (16 %, 20 %, 25 %, 32 %, 40 %, 50 % and 63 % (*w/v))* of a certain sugar was tested. Contaminations at the antennae were immediately removed and rinsed with water. An inter-trial interval of two minutes was used to prevent intrinsic sensitization [24]. It has been shown that the order of sugar water concentration does not influence responsiveness to sucrose [23]. For each sugar and each bee, the positive PER of the concentrations was recorded individually. The sum of responses to all concentrations of a certain sugar constitutes the individual gustatory response score (GRS) of a bee, which is a measure for overall responsiveness [23] [24] [25].

### Genotyping via next generation sequencing (NGS)

Directly after the PER tests, bee’s genomic DNA (gDNA) was isolated as described before [13]. The gDNA samples of putative mutants and the control group were preselected via a hex-labeled PCR and fluorescence length analysis (suppl. Tab. S2). Subsequently, we performed next generation sequencing in multiplex approach (NGS, with GENEWIZ, Leipzig, Germany). The samples were indexed with a tag and amplified with adapter overhangs approach (see suppl. Tab. S2). They were sequenced on an Illumina HiSeq 2500 (2×250bp, Rapid Run). Bioinformatic analysis followed the strategy with a handwritten Perl script as used previously by Değirmenci et al. [13]. De-multiplexing of the samples was carried out via their identifier tags using HMMer v3.2.1 [26]. All reads (forward and reverse) were merged and subsequently quality filtered (maxEE=1, minlen=100) using USEARCH v11 [27]. We identified and counted variants of each sample making use of USEARCH for dereplication and counting, therefore, taking into account both alleles of each individual. In this script, also MUSCLE was used for alignment with the reference and afterwards indel positions were counted. For AmGr1, the alignment was split into segments to cover only the relevant site to account for splice variants at other positions [27] before counting indels. Wild-type alleles were classified as “wt” and in-frame indels with a multiple of 3 bps were marked as “if”, resulting in an intact open reading frame and exclusion from our investigation. The label “ns” was used for “nonsense” mutation, since these mutations would result in reading frame shifts (nonsense code), leading to non-functional proteins. We thus followed the proved genotyping approach of Roth et al. [20] and only included animals with a proven homozygous mutant (ns/ns) or homozygous wildtype (wt/wt) genotype.

### Quantification and statistical analysis

For the electrophysiological part of our work, we performed at least two independent experiments (oocytes from different batches). The sample size *n* and statistical details (mean ± standard deviation, SD) are given in the figure legends. Data analysis was performed using the software Igor Pro 8 (waveMetrics, Inc., Lake Oswego, Oregon, USA). For statistical analysis, the software Excel (Microsoft Corp. Redmond, Washington, USA) was used. For the behavioural analysis the GRS of mutant and wild-type bees of each sugar were compared using the Mann-Whitney-U test, since data was not normally distributed.

## Acknowledgments

We thank Markus Thamm for molecular lab expertise and Karin Möller for technical assistance. We thank Katharina Beer, Daniel Rodriquez and Florian Loidolt for honeybee queen caging, egg collection, preparation for injection and help with artificial rearing. We thank Martin Gabel, Lioba Hilsmann, Florian Loidolt and Dirk Ahrens for the beehive maintenance and Stefan Berg for providing the beehives. This project was funded by a grant of the German Research Foundation to R.S. (SCHE 1573/8-1) and by the Volkswagen Foundation to R.S.

## Author Contributions

L.D., F.L.R.F, D.G. and R.S. designed research and drafted the manuscript. D.G. and F.L.R.F. performed biophysical characterization experiments and analysed the data. L.D. and A.V. performed CRISPR/Cas9 experiments and data analysis. C.H. was involved in experimental support and artificial rearing of honeybees. A.K. performed bioinformatics for genotyping. All authors contributed to the final manuscript and approved it.

## Availability of data and materials

Correspondence and requests for materials not included in the supplementary information should be addressed to L.D. and for CRISPR/Cas9, to L.D. for *in vitro* rearing of larvae, to D.G. and F.L.R.F. for characterization and to A.K. for bioinformatics.

## Ethics approval and consent to participate

No ethics approval or consent to participate was required for this study.

## Supplementary Information

For:

**The following supporting information is available for this article:**

**Figure S1:**
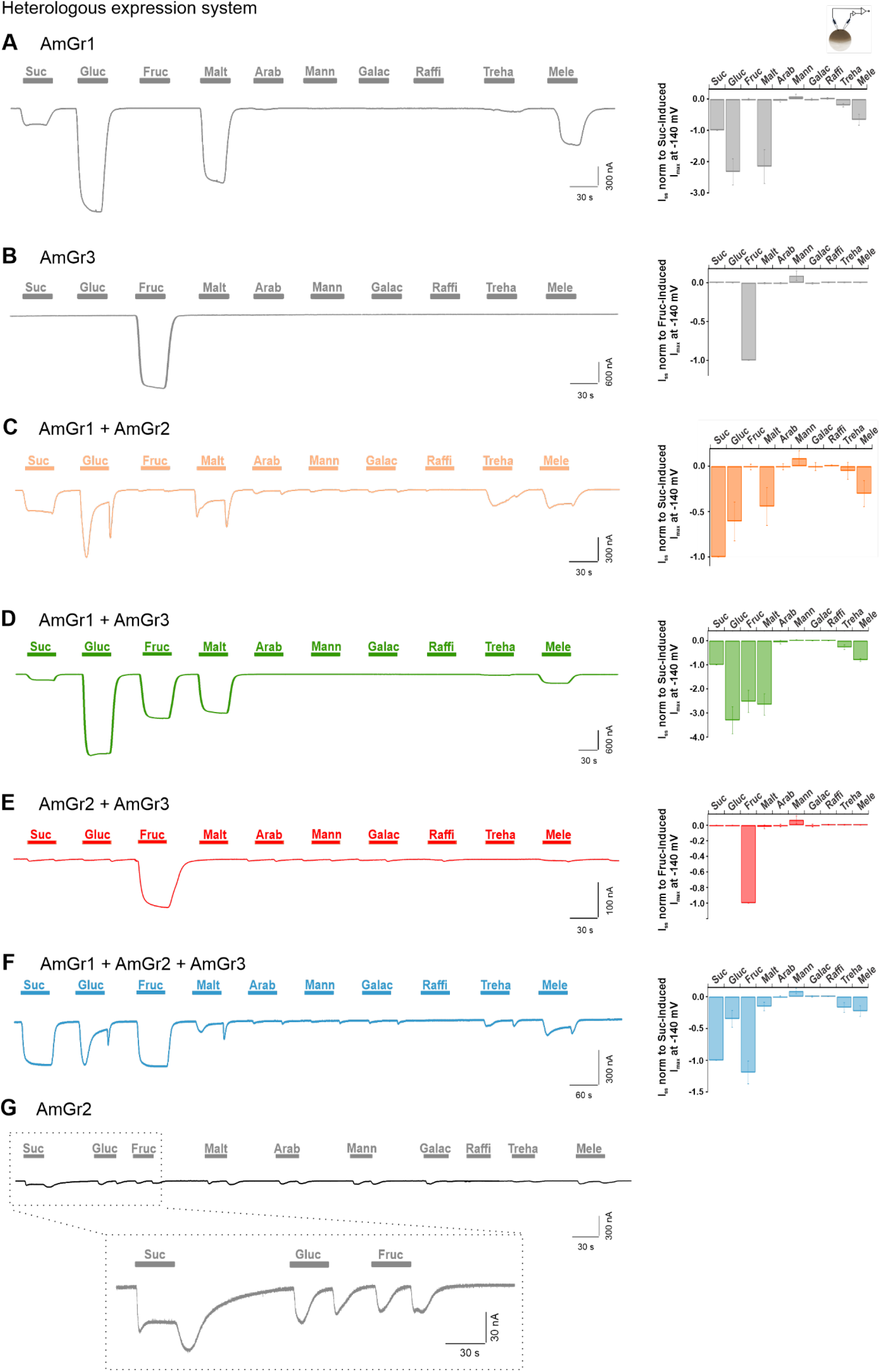
Representative TEVC recordings of sugar-induced currents derived from *Xenopus* oocytes expressing different Gr-ensembles. The gustatory receptors AmGr1-3 have been transiently expressed in *Xenopus* oocytes either alone or in different Gr combinations. Oocytes were clamped at a holding membrane potential of −80 mV and tested for sugar response to sucrose (Suc), glucose (Gluc), fructose (Fruc), maltose (Malt), arabinose (Arab), mannose (Mann), galactose (Galac), raffinose (Raffi) and melezitose (Mele) (160 mM each; perfusion indicated by bars). For quantification of sugar specificities, sugar-induced steady-state currents (*I_SS_*) were recorded at a membrane potential of −140 mV and normalized to the currents in either sucrose (A, C, D and F) or fructose (B and E) solution (bar diagrams). **A** AmGr1 (mean of *n* = 16 oocytes ± SD); **B** AmGr3 (mean of *n* = 8 oocytes ± SD); **C** AmGr1 and AmGr2 co-expression (mean of *n* = 13 oocytes ± SD); **D** AmGr1 and AmGr3 co-expression (mean of *n* = 13 oocytes ± SD); **E** AmGr2 and AmGr3 co-expression (mean of *n* = 9 oocytes ± SD); **F** AmGr1, AmGr2 and AmGr3 co-expression (mean of *n* = 10 oocytes ± SD); **G** AmGr2, Inset: magnification of the current trace of AmGr2-expressing oocyte for sucrose, glucose and fructose application.

**Figure S2:**
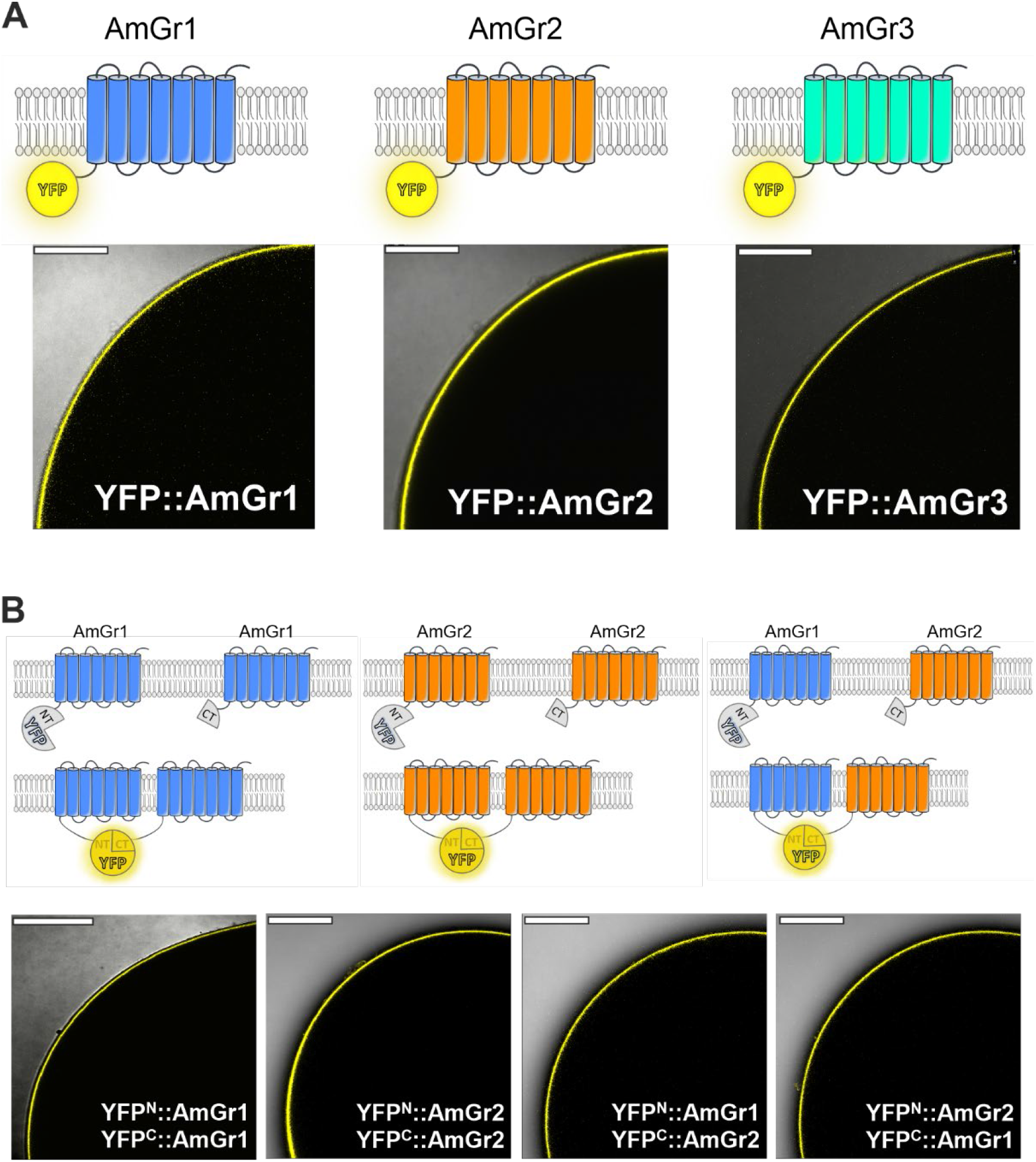
Fluorescence-based studies of *Xenopus* oocytes expressing YFP-tagged AmGr1-3 constructs. Schematic models of YFP-tagged AmGrs and AmGrs tagged with YFP halves for BIFC experiments (A and B, upper panels); Pictures show a quarter of an optical slice of an oocyte, representing an overlay of brightfield and detection of fluorescence (depicted in yellow). Images were taken with a confocal laser scanning microscope (A and B, lower panels). (A) Representative images of gustatory receptors AmGr1-3 tagged with YFP to the N-terminus (YFP::AmGr1; YFP::AmGr2; YFP::AmGr3) transiently expressed in *Xenopus* oocytes. **(B)** Interaction studies of AmGr1 and AmGr2 by bimolecular fluorescence complementation (BIFC). N- and C-terminal YFP halves fused to either AmGr1 or AmGr2 subunits complemented YFP fluorescence when co-expressed in oocytes, indicating physical interaction, i.e. via homomerization of AmGr1 subunits (YFP^N^::AmGr1 + YFP^C^::AmGr1) or AmGr2 subunits (YFP^N^::AmGr2 + YFP^C^::AmGr2). When corresponding YFP halves were fused to AmGr1 and AmGr2 (YFP^N^::AmGr1 + YFP^C^::AmGr2 / YFP^N^::AmGr2 + YFP^C^::AmGr1) co-expression in oocytes led to yellow fluorescence (YFP complementation), indicating physical interaction of AmGr1 and AmGr2 subunits that assemble to heterotetrameric receptors. (Scale bar = 200 μm)

**Figure S3:**
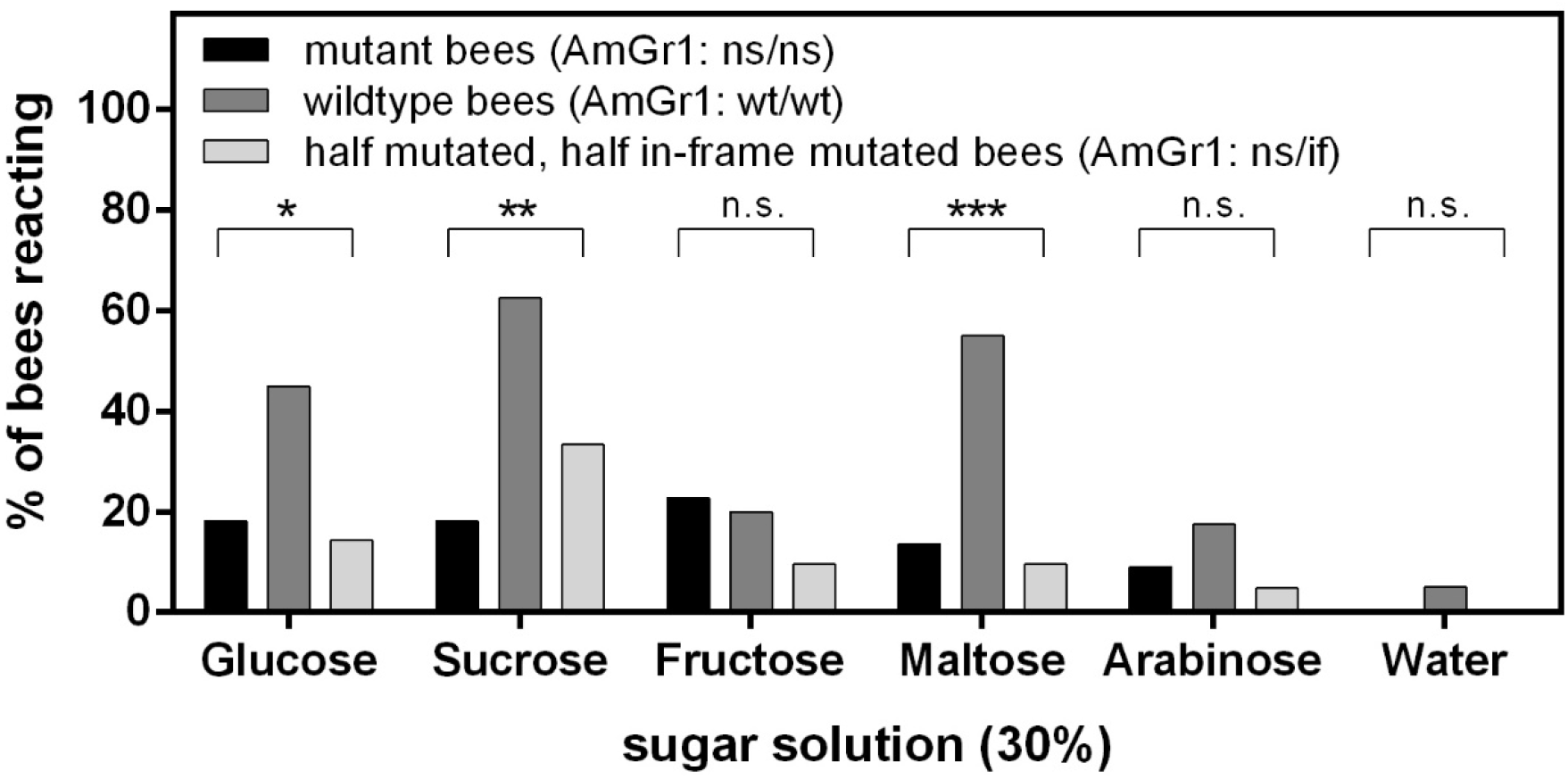
Behavioural PER test with AmGr1 mutants (ns/ns), half-mutated and half in-frame-mutated bees of AmGr1 (ns/if) and wildtype bees (wt/wt) with a 30% solution of several other sugars (glucose, sucrose, fructose, maltose, arabinose) and water. The percentage of bees reacting to the representative sugar (only 30% solution, only one PER-test) is shown. The respective contingency tables of the three groups (N(ns/ns)=22, N(wt/wt)=40 and N(ns/if)= 21, portioned in reacting and not reacting bees) were analysed with a Chi-Square test and showed, that they were significantly different for glucose (Chi-Square test; Chi^2=^8.199;df=2; p=0.0166; *), sucrose (Chi-Square test; Chi^2=^12.5; df=2; p=0.0019; **) and maltose (Chi-Square test; Chi^2=^17.84; df=2; p=0.0001; ***). In these cases, wildtype bees (wt/wt, dark grey middle bars) reacted with a bigger portion than the AmGr1 mutants (ns/ns, black left bars) or the half-mutated and half in-frame-mutated bees (ns/if, light grey right bars). For the sugars fructose (Chi-Square test; Chi^2=^1.459; df=2; p=0.4822; n.s.) and arabinose (Chi-Square test; Chi^2=^2.356; df=2; p=0.3079, n.s.) and for water (Chi-Square test; Chi^2=^2.203; df=2; p=0.3324; n.s.) there were no significant differences in the three groups. These findings are consistent with the results of the TEVC recordings of additional sugars (Figure S1).

**Figure S4:**
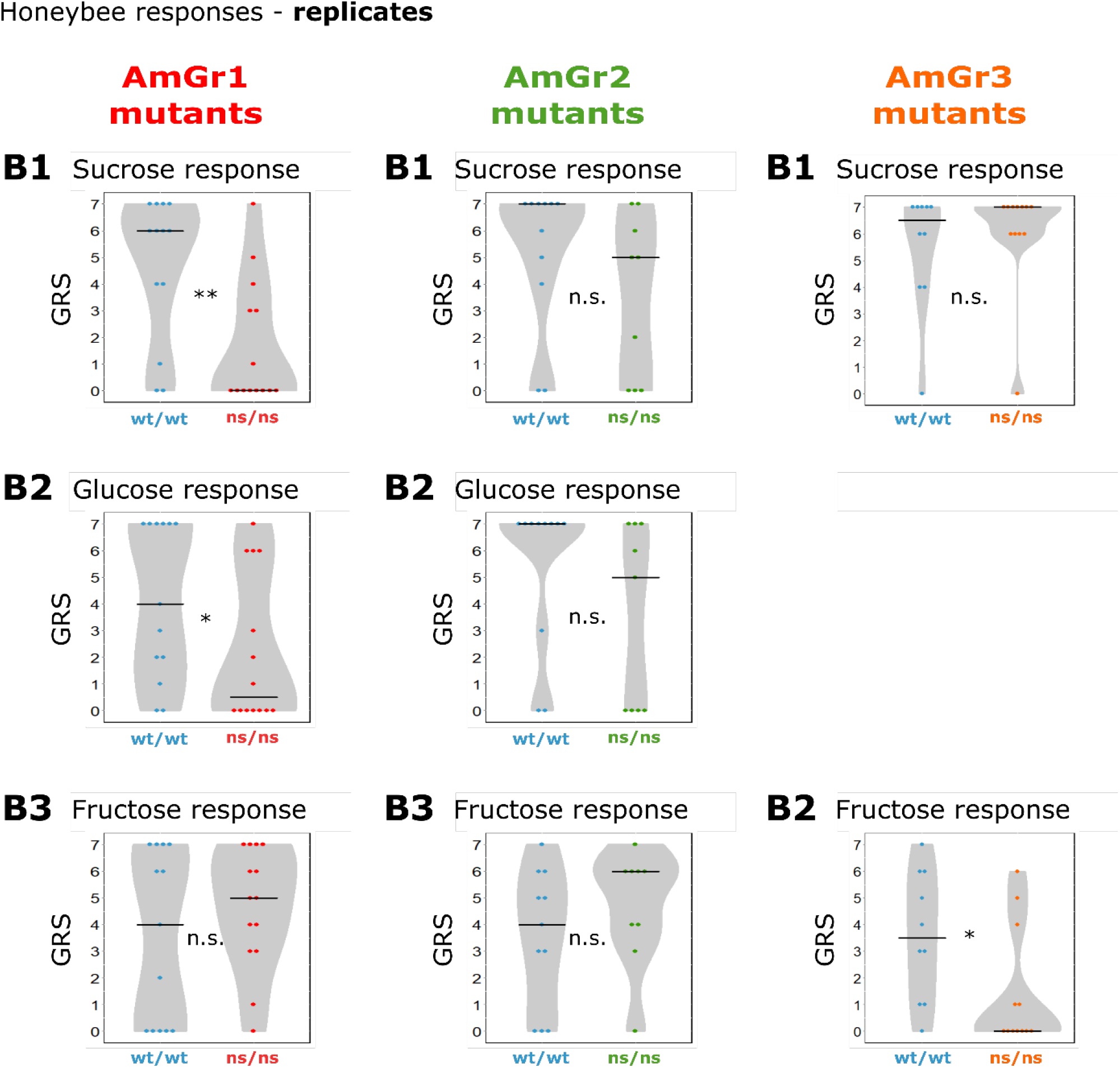
Replicates of behavioural evaluation through proboscis extension response (PER, *in vivo)* according to the section B of each figure (Fig. 1-3). AmGr1 mutant bees (ns/ns; N= 14, red) are significantly less responsive towards sucrose (B1) compared to wild-type bees (wt/wt; N=13, blue) (AmGr1 mutants: B1; Mann-Whitney-U, ns/ns vs. wt/wt, p=0.0051, **). Glucose responsiveness (B2) of wild-type bees is significantly higher than those of AmGr1 mutants (AmGr1 mutants: B2; Mann-Whitney-U, ns/ns vs. wt/wt, p=0.0412, *). Both groups do not differ when comparing their responsiveness towards fructose (AmGr1 mutants: B3; Mann-Whitney-U, ns/ns vs. wt/wt, p=0.4528, n.s.). AmGr2 mutant bees (ns/ns; N=9, green) do not differ in their responsiveness towards sucrose (AmGr2 mutants: B1; Mann-Whitney-U, ns/ns vs. wt/wt, p=0.1940, n.s.), glucose (AmGr2 mutants: B1; Mann-Whitney-U, ns/ns vs. wt/wt, p=0.1284, n.s.) or fructose (AmGr2 mutants: B1; Mann-Whitney-U, ns/ns vs. wt/wt, p=0.3099) when compared with wild-type bees (wt/wt; N=11, blue). AmGr3 mutant bees (ns/ns; N= 12, orange) do not differ in their responsiveness towards sucrose (B1) compared to wild-type bees (wt/wt; N=10, blue) (AmGr3 mutants: B1; Mann-Whitney-U, ns/ns vs. wt/wt, p=0.5383, n.s.). AmGr3 mutants are significantly less responsive to fructose than wildtype bees (AmGr3 mutants: B2; Mann-Whitney-U, ns/ns vs. wt/wt, p=0.0266, *). Both published previously in Degirmenci et al. 2020.

**Table S1:**
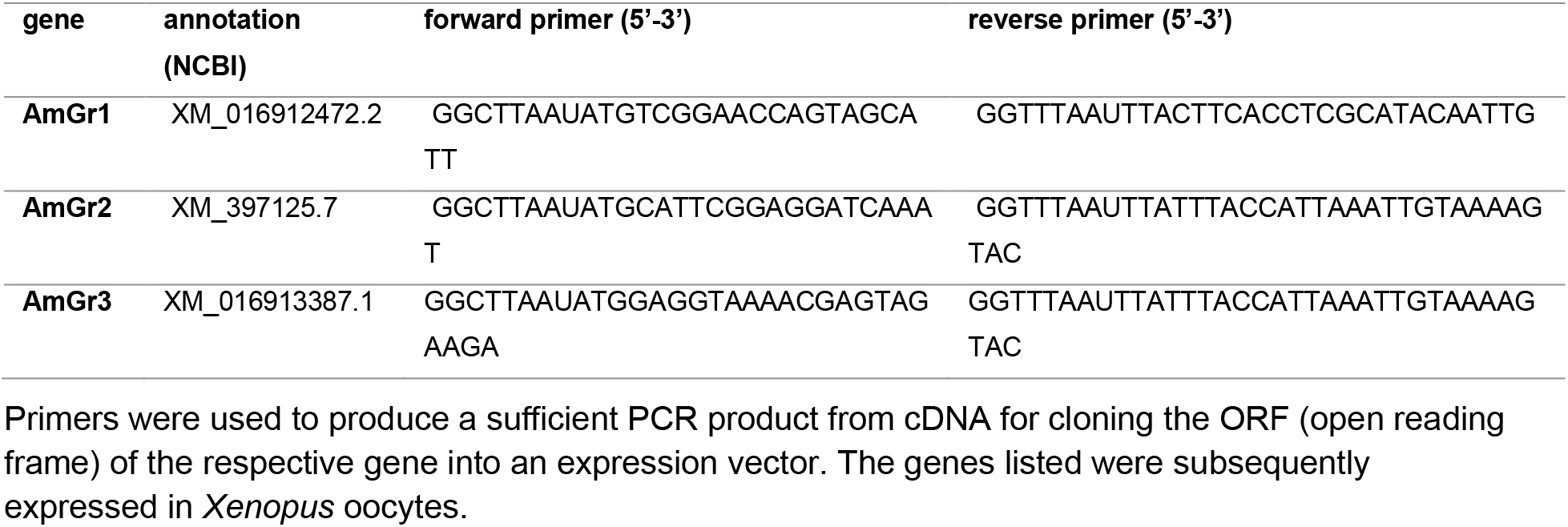
Annotations and primers used for cloning the respective AmGr genes into expression vectors.

**Table S2:**
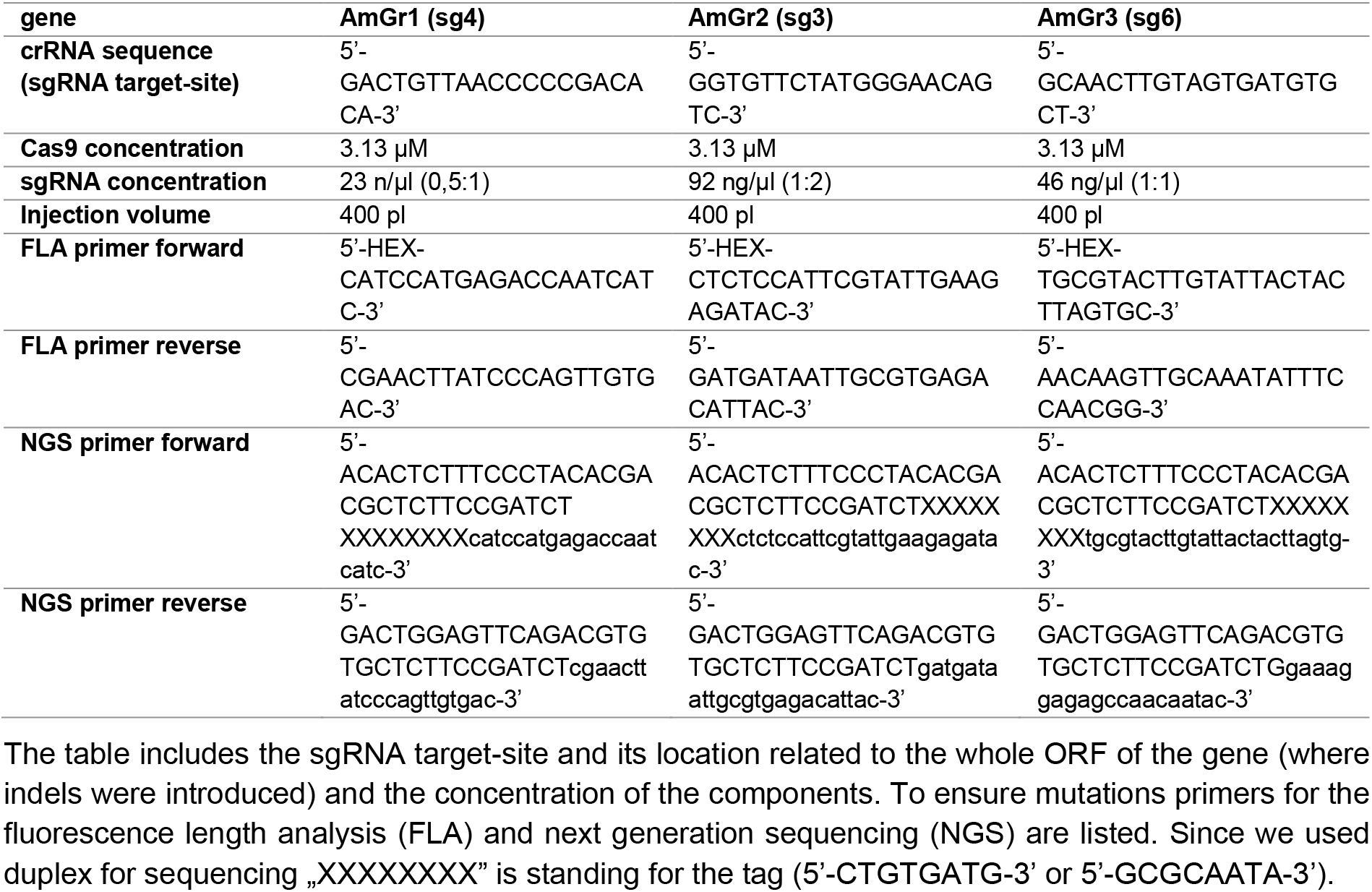
Sequence information for performing CRISRP/Cas9 in the genes of the sugar receptors AmGr1, AmGr2 and AmGr3.

